# clinker & clustermap.js: Automatic generation of gene cluster comparison figures

**DOI:** 10.1101/2020.11.08.370650

**Authors:** Cameron L.M. Gilchrist, Yit-Heng Chooi

## Abstract

**Summary:** Genes involved in biological pathways are often collocalised in gene clusters, the comparison of which can give valuable insights into their function and evolutionary history. However, comparison and visualisation of gene cluster homology is a tedious process, particularly when many clusters are being compared. Here, we present `clinker`, a Python based tool, and `clustermap.js`, a companion JavaScript visualisation library, which used together can automatically generate accurate, interactive, publication-quality gene cluster comparison figures directly from sequence files.

**Availability and Implementation:** Source code and documentation for `clinker` and `clustermap.js` is available on GitHub (github.com/gamcil/clinker and github.com/gamcil/clustermap.js, respectively) under the MIT license. `clinker` can be installed directly from the Python Package Index via `pip`.

## Introduction

Genes involved in biological pathways are often collocalised in the genome in gene clusters, the study of which is important given their potential to encode traits such as the biosynthesis of bioactive small molecules, virulence and drug resistance. Gene clusters are typically conserved to some extent across taxa, and comparative analysis can reveal significant insights into their evolution and potential differences in the pathways they encode. For instance, the conservation (or lack thereof) of gene clusters encoding the biosynthesis of secondary metabolites can guide the search for new drug compounds and bioactivities (Gilchrist *et al.*, 2018, Ziemert *et al.*, 2016).

Comparative analysis of large genomic datasets has become increasingly common in the post-genomics era, with visualisations of gene cluster conservation across species often featured prominently in such work (de Vries *et al.*, 2017, Nielsen *et al.*, 2017, Doroghazi and Metcalf, 2013). This is a tedious process, and as such has driven the development of numerous tools (e.g. EasyFig (Sullivan *et al.*, 2011), Artemis Comparison Tool (Carver *et al.*, 2005), GeneSpy (Garcia *et al.*, 2019), Gene Graphics (Harrison *et al.*, 2018)) and libraries (e.g. GenomeDiagram (Pritchard *et al.*, 2006), genoPlotR (Guy *et al.*, 2010)). However, these tools have drawbacks: most have limited interactivity, with clusters unable to be repositioned, reordered or resized without changing input files; visualisation options are inflexible, if not fixed; some require sequence comparison files to be generated externally using tools such as BLAST (Camacho *et al.*, 2009), complicating the process; and libraries require some level of programming knowledge, making them inaccessible to the average biologist. Thus in many cases, researchers choose either to forgo specialist tools entirely, creating figures manually in other software (e.g. Microsoft PowerPoint, Adobe Illustrator), or directly use the output generated by analysis tools such as MultiGeneBlast (Medema *et al.*, 2013), cblaster (Gilchrist, 2020), or antiSMASH (Blin *et al.*, 2019). This quickly becomes a daunting prospect when many clusters are to be compared. There is therefore a need for tools which produce clear, publication-quality visualisations, are more flexible and intuitive than existing tools, and require no significant effort on the part of the user other than providing sequence data.

Here we describe `clinker`, a Python-based gene cluster comparison pipeline, and `clustermap.js`, a companion visualisation JavaScript library, which can automatically generate interactive, to-scale gene cluster comparison figures directly from sequence files.

## Gene cluster alignments using clinker

The `clinker` workflow is detailed in Figure 1a. `clinker` accepts GenBank files as input; multi-record GenBank files are treated as gene clusters with multiple loci. Amino acid translations of genes in each cluster are extracted and aligned using the BioPython package (Cock *et al.*, 2009). By default, global alignments are performed using the BLOSUM62 subsitution matrix, a gap open penalty of −10 and extension penalty of −0.5, however parameters can be changed when directly using the clinker Python API. Any alignments not reaching the user-defined sequence identity threshold are discarded.

**Figure 1:**
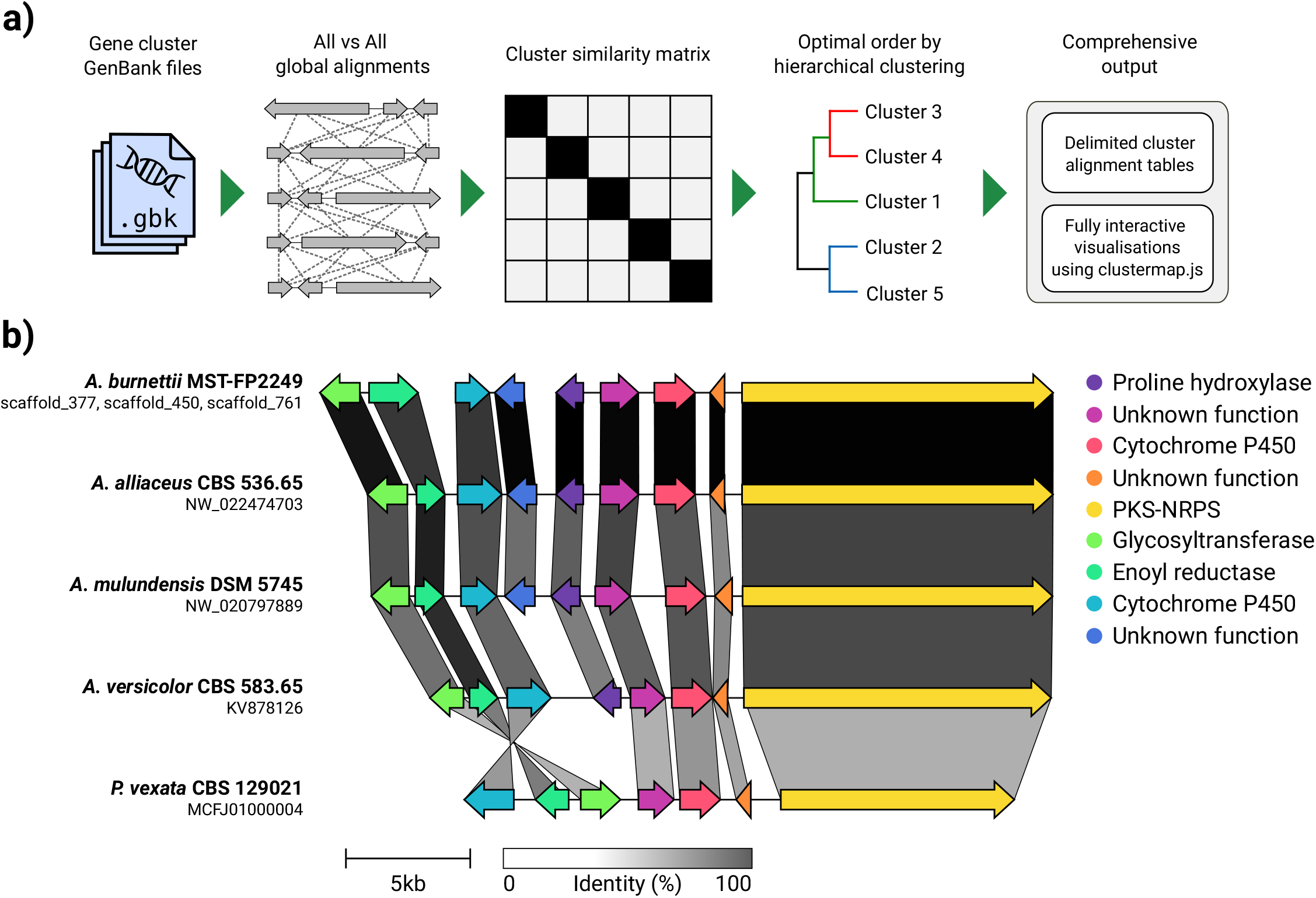
The `clinker` pipeline (a) and visualisation of the burnettramic acid biosynthetic gene cluster (Li *et al.*, 2019) and homologues generated by `clinker` using `clustermap.js` (b). The gene cluster comparison figure in (b) is presented exactly as was achieved within the `clustermap.js` visualisation, with only minor further editing of text labels done in external software (italicising species names). The static HTML file used to generate (b) is provided in the supplementary information.

Optimal ordering of clusters for visualisation is determined through hierarchical clustering. First, a cluster similarity score is calculated for every pair of input clusters. clinker uses a modified version of the formula implemented by Medema *et al.* (2013) in MultiGeneBlast, which incorporates both sequence homology and syntenic conservation. Briefly, the similarity of two clusters is given by *S* = *h* + *i* · *s*, where *h* is the cumulative sequence identity of homologous sequences in each cluster, s is the total number of shared contiguous sequence pairs, and *i* is a weighting factor which determines the weight of synteny compared to sequence homology (0.5 by default). The resulting similarity matrix is then hierarchically clustered using the Ward variance minimization algorithm as implemented in the SciPy package (Virtanen *et al.*, 2020). The leaves of the dendrogram generated in this step are used as the default order of clusters in the clustermap.js visualisation.

## Interactive visualisations using clustermap.js

Cluster alignments generated using `clinker` are visualised using `clustermap.js` (Figure 1b). Though designed in conjunction with `clinker, clustermap.js` is a standalone library and can take data generated elsewhere as long as it abides by the correct schema.

Clusters are drawn to scale based on a user-definable scaling factor (15 pixels per 1000 base pair by default). Clusters can be renamed by clicking their labels and reordered by dragging them, enabling comparisons between clusters outside of the computed optimal ordering.

`clustermap.js` is capable of displaying multi-locus clusters. For example, the biosynthetic gene cluster for the burnettramic acids (Li *et al.*, 2019) is split over three contigs due to fragmented genome assembly, but is readily visualised by our tool (top-most cluster in Figure 1b). Hovering over a locus displays a grey box with handles at both ends; loci can be freely repositioned by clicking and dragging the box, inverted by double clicking it, or resized (hiding genes) by dragging the handles. Any changes in the visible area of specific loci is reflected by the coordinates in the cluster label. Genes are drawn as arrows, whose shape is easily altered using sliders for body height, tip height and length. The visualisation can be anchored around a specific gene by clicking on it: `clustermap.js` will identify the closest homologous genes in other clusters and automatically reposition them to align with the clicked gene. Gene labels can be resized, repositioned or hidden entirely.

Links are drawn between homologous genes on neighbouring clusters and are shaded based on sequence identity (0% white, 100% black). By default, all links above the identity threshold in `clinker` are drawn; this threshold can be raised within the `clustermap.js` visualisation to dynamically hide links. Alternatively, one can choose to show only the highest scoring links between clusters, which is useful when multiple similar genes exist within a cluster, resulting in many overlapping links. Groups of homologous genes are established via single linkage of gene links. Each homology group is assigned a unique colour, which is used as the fill of both the genes in the group and its corresponding entry in the figure legend.

The scale bar, colour bar and homology group legend are embedded in the figure, and are automatically repositioned based on cluster and locus placement. The length of the scale bar can be changed by clicking on its text. Likewise, legend entries can be renamed by clicking their text.

Figures generated by `clinker` and `clustermap.js` are easily exported as scalable vector graphics (SVG) files that can be further modified in vector image software.

`clinker` and `clustermap.js` can serve as a useful extension of gene cluster search tools such as MultiGeneBlast (Medema *et al.*, 2013) or cblaster (Gilchrist, 2020).

In conclusion, `clinker` and `clustermap.js` enable the easy creation of publication quality gene cluster comparison figures, and can help facilitate and accelerate large-scale comparative genomic analysis, visualisation and presentation of gene clusters.

## Software Implementation and Availability

`clinker` is implemented in Python 3 and can be freely installed from the Python Package Index (pypi.org/project/clinker). `clustermap.js` is implemented in JavaScript and is built on top of the D3 visualisation library (Bostock *et al.*, 2011). Source code and documentation for both tools is freely available on GitHub (github.com/gamcil/clinker and github.com/gamcil/clustermap.js).

## Acknowledgements

C.L.M.G is supported by an Australian Government Research Training Project scholarship. Y-H.C is supported by an Australian Research Council Future Fellowship (FT160100233).

